# Nerve Injury-Induced Protein 2 preserves lysosomal membrane integrity to suppress ferroptosis

**DOI:** 10.64898/2026.02.09.704867

**Authors:** Jin Zhang, Miranda Bustamante, Yang Shi, Ken-ichi Nakajima, Xinbin Chen

## Abstract

Nerve injury-induced protein 1 (NINJ1), a cell adhesion molecule, is oligomerized during lytic cell death and mediates plasma membrane rupture to release large intracellular molecules that propagate the inflammatory response. We and others previously showed that NINJ2, a close relative of NINJ1, does not promote plasma membrane rupture to spread inflammation. Here, we identify that NINJ2 is necessary for the lysosome membrane integrity to protect cells from ferroptosis. Specifically, we found that NINJ2 localizes to lysosomes and interacts with LAMP1, an anchor glycoprotein of the lysosome membranes and a sensor of stressed lysosomes. We also found that loss of NINJ2 exacerbates lysosomal membrane permeabilization (LMP), which allows for selective leakage of lysosomal contents, such as labile iron, into the cytosol. Accordingly, loss of NINJ2 elevates cellular labile iron accumulation and decreases expression of ferritins, the primary intracellular iron storage protein complexes. Mechanistically, we found that loss of NINJ2 promotes ferritin FTH degradation in lysosomes, which can be reversed by knockdown of LAMP1. Moreover, we found that loss of NINJ2 sensitizes cells to ferroptosis induced by RSL3 and Erastin, consistent with a recent study that loss of *Ninj2* predisposes mice to chronic inflammation. Together, these findings uncover a previously unrecognized activity of NINJ2 from lysosome homeostasis to ferroptosis, which can be explored as a cancer therapeutic strategy especially considering that NINJ2 and ferritins are found to be overexpressed and positively associated with iron-addicted cancers.

## Introduction

Ninjurin 2 (Nerve Injury-Induced Protein 2; NINJ2), along with NINJ1, belongs to the Ninjurin family of homophilic cell-surface adhesion molecules [1, 2]. These proteins were originally identified as being upregulated in Schwann cells and dorsal root ganglion neurons following peripheral nerve injury. Subsequent studies showed that NINJ1 and NINJ2 promote neurite outgrowth and facilitate interactions between Schwann cells and regenerating axons, thereby contributing to effective nerve repair [1–3]. Structural analysis indicated that NINJ1 and NINJ2 share approximately 52%–55% amino acid sequence identity and 67% sequence similarity. They are a two-pass transmembrane protein composed of an extracellular N-terminal domain, two conserved hydrophobic transmembrane domains, and a C-terminal extracellular domain [4]. Despite high sequence homology, recent studies showed that NINJ1 and NINJ2 exert divergent functions potentially due to their structural variations. For instance, NINJ1 mediates plasma membrane rupture by polymerizing into straight, amphipathic filaments that disrupt membrane integrity [5–7]. In contrast, NINJ2 assembles into curved filaments that are unable to permeabilize the membrane and does not promote plasma membrane rupture [7]. These data indicated that NINJ1 and NINJ2 exhibit both overlapping and distinct functions across diverse biological processes.

Recent studies have shown that NINJ2 is a multifaced protein with profound biological functions. NINJ2 is found to be highly expressed in immune-related tissues, including bone marrow, lymph nodes, spleen, and thymus [8, 9], suggesting a role in immune regulation. Indeed, studies have shown that NINJ2 acts as a pro-inflammatory mediator by physically interacting with TLR4, activating the NF-κB pathway to promote the expression of inflammatory markers like IL-6 and TNF-α [10–12]. Additionally, we found that loss of NINJ2 leads to activation of NLRP3 inflammasome and subsequently, promotes pyroptosis, a pro-inflammatory programmed cell death that leads to the dysregulated release of cytokines like IL-1β and IL-18 [9]. Furthermore, we recently identified NINJ2 as a transcriptional target of the tumor suppressor p53 and that NINJ2, in turn, represses p53 mRNA translation, implicating its role in tumorigenesis [13]. Finally, *NINJ2* gene polymorphisms are found to be associated with increased risk of ischemic stroke, coronary artery disease, multiple sclerosis (MS), and Alzheimer’s disease [14–17], suggesting a role in vascular and neuroinflammatory pathologies. Together, these data indicate that NINJ2 participates in a wide range of physiological and pathological processes, yet the molecular mechanisms underlying its multifunctional roles remain poorly understood.

Lysosomes are central regulators of cellular homeostasis, coordinating protein degradation, autophagy, and metabolic signaling [18]. Lysosomal membrane permeabilization (LMP) is defined by the formation of ultrastructurally-undetectable, tiny pores at the lysosomal membrane and thereby causes selective releases of lysosomal contents into the cytosol [19, 20]. Recently, it was found that low-grade LMP does not always cause cell death but can be repaired by ESCRT complex [21]. It was also found that LMP triggers the activation of NLRP3 inflammasome and subsequently, elicits immune response [22, 23]. We previously reported that NINJ2 deficiency leads to NLRP3 inflammasome activation [9]. However, it was not certain whether this activation was linked to lysosomal damage. To address this, we examined the role of NINJ2 in regulating lysosomal morphology and function. We found that NINJ2 localizes to lysosomes and loss of NINJ2 exacerbates LMP. We also found that the LMP mediated by NINJ2-deficiency results in increased levels of labile iron along with decreased expression of ferritin. We further demonstrated that NINJ2-deficiency promotes degradation of Ferritin and subsequently, enhances ferroptosis, a regulated form of cell death driven by iron-dependent lipid peroxidation. Together, these findings reveal a previously unrecognized function of NINJ2 in linking lysosomal homeostasis to ferroptosis.

## Results

### Ninj2 protein localizes to Lysosomes and interacts with LAMP1

To determine whether NINJ2 is involved in modulating lysosomal activity, we first examined whether NINJ2 localizes to lysosomes. To this end, MCF7 cells expressing Flag-tagged NINJ2 were stained with LysoTracker to label lysosomes and with an anti-Flag antibody to detect NINJ2. Confocal microscopy revealed substantial co-localization of NINJ2 with LysoTracker-positive vesicles, indicating that some NINJ2 proteins localize to lysosomes (Fig. 1A). To further test this, MCF7 cells expressing Flag-tagged NINJ2 were co-stained with antibodies against NINJ2 and lysosome-associated membrane protein 1 (LAMP1), a well-established lysosomal marker [24–26]. Consistent with the data from LysoTracker staining (Fig. 1A), we found that NINJ2 was co-localized with LAMP1 in lysosomes (Fig. 1B), confirming that NINJ2 localizes to lysosomes. Next, we examined whether NINJ2 physically interacts with LAMP1 by performing reciprocal immunoprecipitation assays using 293T cells expressing Flag-tagged NINJ2. We found that endogenous LAMP1 was detectable in NINJ2-immunocomplex when immunoprecipitated with anti-Flag antibody (Fig. 1C). Conversely, Flag-tagged NINJ2 was readily detected in LAMP1-immunocomplexes when immunoprecipitated with anti-LAMP1 antibody (Fig. 1D). To verify that NINJ2 interacts with LAMP1, we performed proximity ligation assay (PLA) [27], which detects protein-protein interactions at a subcellular level. Briefly, MCF7 cells expressing Flag-tagged NINJ2 were incubated with anti-Flag and/or anti-LAMP1 antibodies. PLA signals were visualized as discrete fluorescent puncta. We observed robust PLA signals in cells stained with both antibodies, indicating close interaction between NINJ2 and LAMP1 (Fig. 1E). In contrast, no PLA signals were detected in control conditions in which cells were incubated without any primary antibody or with only one antibody (Fig. 1E).

**Figure 1.**
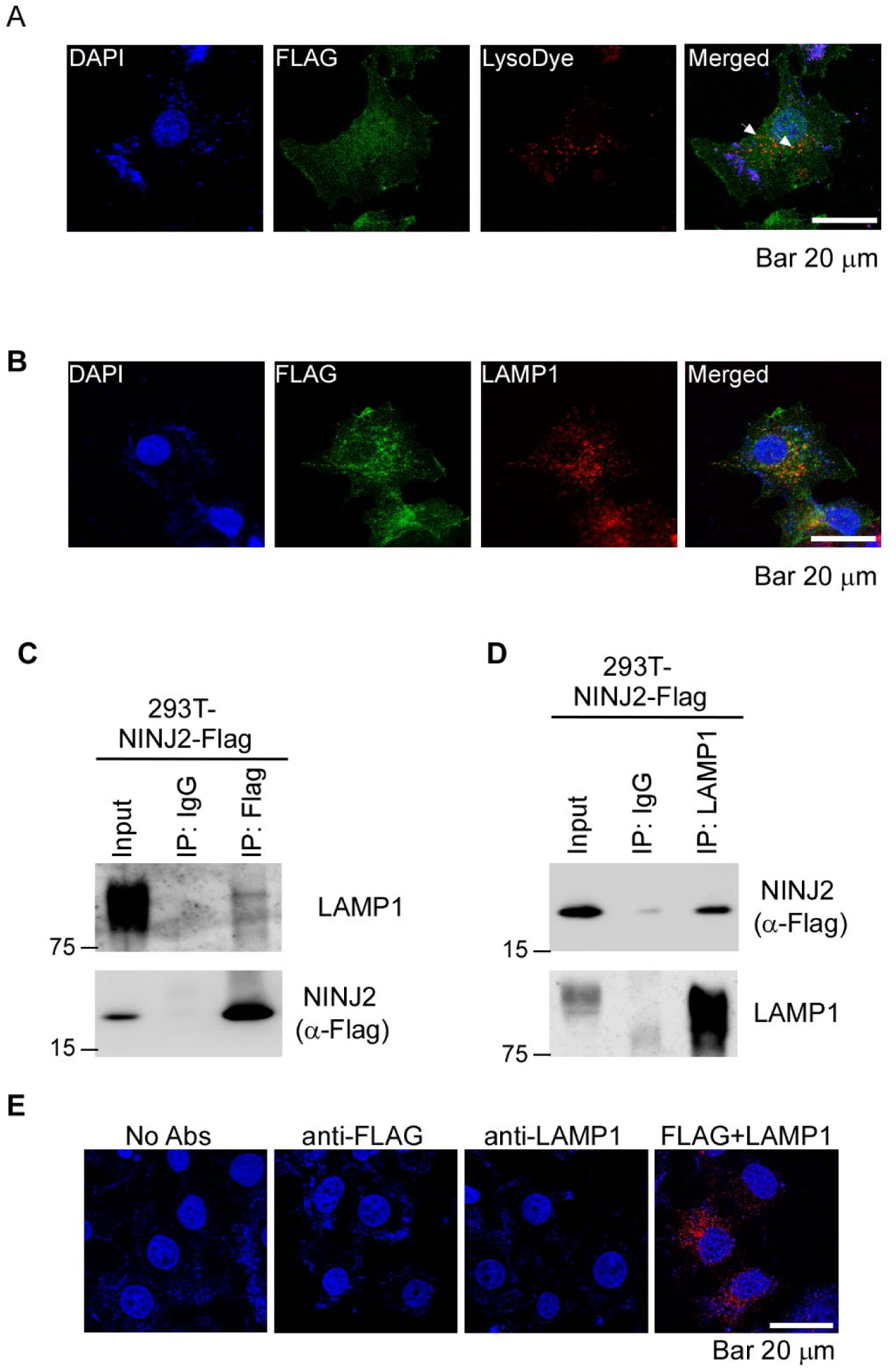
NINJ2 protein localizes to lysosomes and interacts with LAMP1. (**A**) MCF7 cells were transiently transfected with a plasmid expressing Flag-tagged NINJ2, followed by immunostaining with, DAPI, lysoDye and Flag. The arrow indicates co-staining of NINJ2 and LysoDye. (**B**) MCF7 cells were transiently transfected with a plasmid expressing Flag-tagged NINJ2, followed by immunostaining with anti-Flag and anti-LAMP1. (**C-D**) 293T cells were transiently transfected with Flag-tagged NINJ2 plasmid for 24 hours, followed by immunoprecipitation with IgG, anti-Flag (C) or anti-LAMP1 (D). The immunocomplex were examined by western blot analysis with Flag or LAMP1 antibody. (**E**) MCF7 cells were transfected with Flag-tagged NINJ2 antibody, followed by PLA assay as described in “Material and Methods”. The positive PLA signal is shown in red puncta.

### Loss of NINJ2 leads to enhanced lysosomal membrane permeability along with increased expression of LAMP1

The localization of NINJ2 in lysosomes prompted us to determine whether NINJ2 regulates lysosomal function. To this end, isogenic control and NINJ2-KO MCF7 cells were mock-treated or treated with L-leucyl-L-leucine methyl ester (LLOMe), a well-established inducer of lysosomal membrane permeabilization (LMP) [28, 29], followed by immunostaining with Galectin-3 and LAMP1. We would like to note that in response to lysosomal damage, Galectin-3 rapidly translocalizes to lysosomal membranes where it participate in lysosomal repair and removal [30, 31]. Thus, colocalization of Galectin-3 with LAMP1 serves as a key indicator of LMP. We found that in the absence of LLOMe, little colocalization of Galectin-3 and LAMP1 was observed in isogenic control and NINJ2-KO MCF7 cells (Fig. 2A, top two panels), indicating minimal lysosomal membrane damage (Fig. 2A, compare top two “MERGE” panels). Upon treatment with LLOMe, isogenic control cells exhibited moderate co-staining of Galectin-3 and LAMP1 (Fig. 2A, Iso. Ctrl+ LLOMe panel). Strikingly, the recruitment of Galectin-3 to LAMP1-positive lysosomes was markedly enhanced in NINJ2-KO MCF7 cells, suggesting that NINJ2 deficiency exacerbates LLOMe-induced lysosomal membrane permeabilization (Fig. 2A, NINJ2-KO+LLOMe panel). In addition to elevated LMP mediated by NINJ2-deficiency, we observed that LAMP1 staining was enhanced in NINJ2-KO MCF7 cells compared to isogenic controls regardless of LLOMe treatment (Fig. 2A, LAMP1 panels), suggesting that NINJ2-deficiency alters LAMP1 expression. To verify this, isogenic control and NINJ2-KO Molt4 cells were mock-treated or treated with LLOMe or glucose oxidase (GO), another LMP inducer [32], followed by measurement of LAMP1 expression. We found that loss of NINJ2 resulted in a marked increase in LAMP1 expression regardless of LLOMe or GO treatment (Fig. 2B-C, LAMP1 panel, compare lanes 1, 3, and 5 with 2, 4, and 6, respectively). In contrast, LLOMe and GO did not significantly alter LAMP1 expression in these cells. Similarly, loss of NINJ2 led to increased LAMP1 expression in MCF7 cells regardless of LLOMe or GO treatment (Fig. 2D-E). Consistent with this, we also showed that the level of LAMP1 transcripts were increased by loss of NINJ2 regardless of LLOME treatment in both Molt4 and MCF7 cells (Supplemental Figure 1). Together, these data suggest that loss of NINJ2 leads to low-grade lysosomal membrane damage accompanied by elevated LAMP1 expression, which in turn exacerbates LMP.

**Figure 2.**
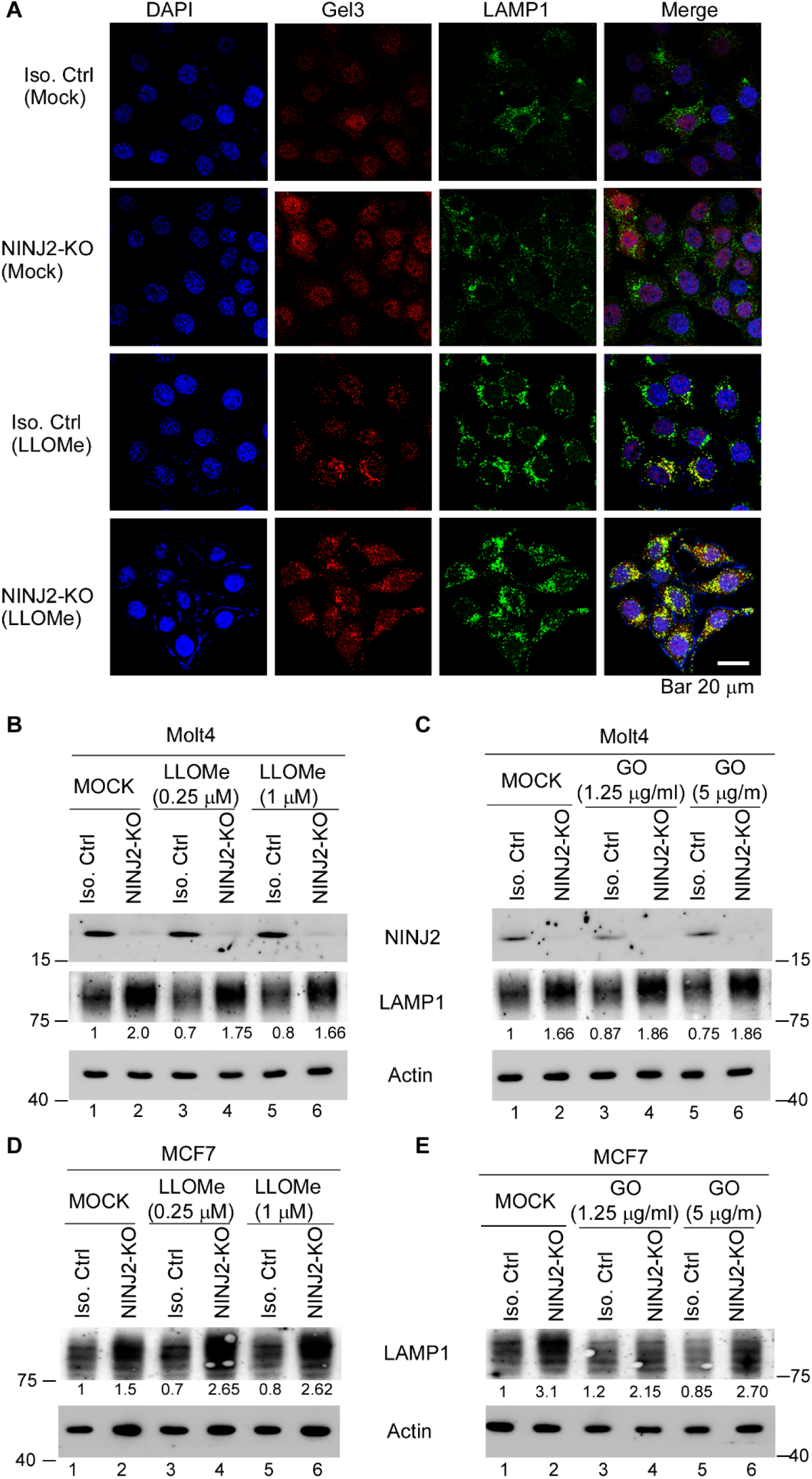
Loss of NINJ2 leads to enhanced LAMP1 expression and lysosomal membrane permeability. **(A)** Isogenic control and NINJ2-KO MCF7 cells were mock-treated or treated with LLOMe (1μM) for 5 hours, followed by immunostaining with antibodies against Galectin 3 and LAMP1. Scale bar: 20μM. (**B-C**) Isogenic control and NINJ2-KO Molt4 cells were mock-treated or treated with LLOMe (B) or GO (C) for 5 hours. The cell lysates were subjected to western blot analysis using antibodies against NINJ2, LAMP1 and actin. The relative protein level of LAMP1 in control cells were arbitrarily set as 1.0 and the relative fold change was shown in below each lane. (D-E) Isogenic control and NINJ2-KO MCF7 cells were mock-treated or treated with LLOMe (D) or GO (E) for 5 hours, followed by western blot analysis to detect expression of LAMP1 and actin. The relative protein level of LAMP1 in control cells were arbitrarily set as 1.0 and the relative fold change was shown in below each lane.

### Loss of NINJ2 elevates intracellular labile iron level and inhibits ferritin expression

LMP is known to release lysosomal contents, such as redox-active Fe²⁺, from lysosomes into the cytosol, thereby expanding the labile iron pool and promoting lipid peroxidation [33]. Thus, we measured the level of labile iron levels in isogenic control and NINJ2-KO Molt4 and MCF7 cells treated with Ferric Ammonium Citrate (FAC), an iron source. We found that the level of intracellular labile iron was markedly increased by loss of NINJ2 in both Molt4 and MCF7 cells (Fig. 3A-B). To verify that NINJ2-deficiency alters iron homeostasis, we measured the expression of ferritin heavy chain (FTH) and ferritin light chain (FTL) in isogenic control and NINJ2-KO Molt4 cells treated with or without FAC. FTH and FTL are oligomerized to form ferritins, the primary intracellular iron storage protein complexes [34, 35]. FTH exerts ferroxidase activity by catalyzing Fe²⁺ to Fe³⁺, whereas FTL helps promote iron nucleation and mineralization for long-term storage. In response to FAC treatment, both FTL and FTH protein were elevated as expected (Fig. 3C-D, FTL and FTH panels, compare lane 1 with 3). Importantly, loss of NINJ2 markedly decreased expression of FTL and FTH (Fig. 3C-D, compare lane 1 and 3 with lane 2 and 4, respectively), suggesting that NINJ2 is required for FTH and FTL expression. Additionally, we found that loss of NINJ2 markedly reduced FTH and FTL expression in MCF7 cells regardless of FAC treatment (Fig. 3E-F, compare lane 1 and 3 with lane 2 and 4, respectively).

**Figure 3.**
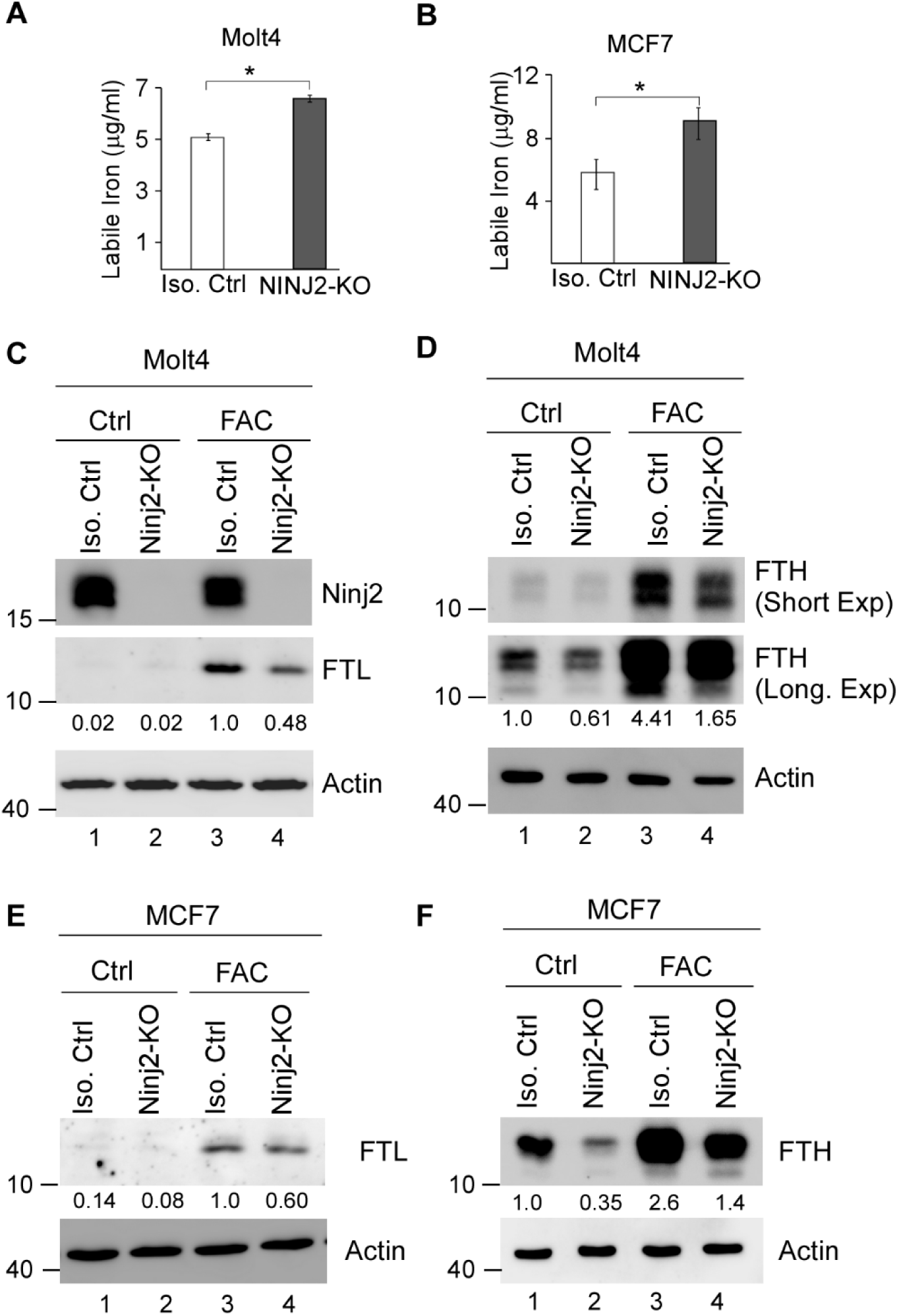
Loss of NINJ2 leads to increased level of labile iron and reduced expression of ferritin. (**A-B**) Isogenic control and NINJ2-KO Molt4 (A) and MCF7 (B) cells were treated with FAC (30mg/ml) for 24 hours. Cell lysates were collected and the level of labile iron was measured using QuantiChrom^TM^ assay kit. * Indicated p<0.05 by student’s test. (**C-D**) Isogenic control and NINJ2-KO Molt4 cells were mock-treated or treated with FAC (30mg/ml) for 16 hours, followed by western blot analysis to measure the level of NINJ2 (C), FTL (C), FTH (D) and actin (C-D). The relative protein level of FTL (C) and FTH (D) in control cells were arbitrarily set as 1.0 and the relative fold change was shown in below each lane. (**E-F**) Isogenic control and NINJ2-KO MCF7 cells were mock-treated or treated with FAC (30mg/ml) for 16 hours, followed by western blot analysis to measure the level of FTL(E), FTH (F) and actin (E-F). The relative protein level of FTL (E) and FTH (F) in control cells were arbitrarily set as 1.0 and the relative fold change was shown in below each lane.

### Loss of NINJ2 promotes ferritin degradation

To uncover the mechanism by which NINJ2 regulates Ferritin expression, we first examined the level of FTH transcripts and found it not to be altered by loss of NINJ2 (Fig. 4A). Next, the half-life of FTH protein was measured in Isogenic control and NINJ2-KO MCF7 cells treated with cycloheximide for various times. We found that the half-life of FTH protein decreased from 23.1h in isogenic control cells to 11.95h in NINJ2-KO cells (Fig. 4B-C), indicating that NINJ2 is required for maintaining FTH protein stability. Since FTH is known to be degraded primarily in lysosomes via NCOA4 [36, 37], we then asked whether LAMP1 plays a role in FTH expression decreased by loss of NINJ2. To this end, two siRNA against LAMP1 were designed and transiently transfected into isogenic control and NINJ2-KO MCF7 cells. As expected, the level of LAMP1 protein was diminished upon siRNA transfection (Fig. 4D, LAMP1). Importantly, knockdown of LAMP1 led to increased expression of FTH and FTL in both isogenic control and NINJ2-KO MCF7 cells (Fig. 4D). Together, these data indicated that loss of NINJ2 promotes Ferritin turnover in lysosomes, which can be reverted by knockdown of LAMP1.

**Figure 4.**
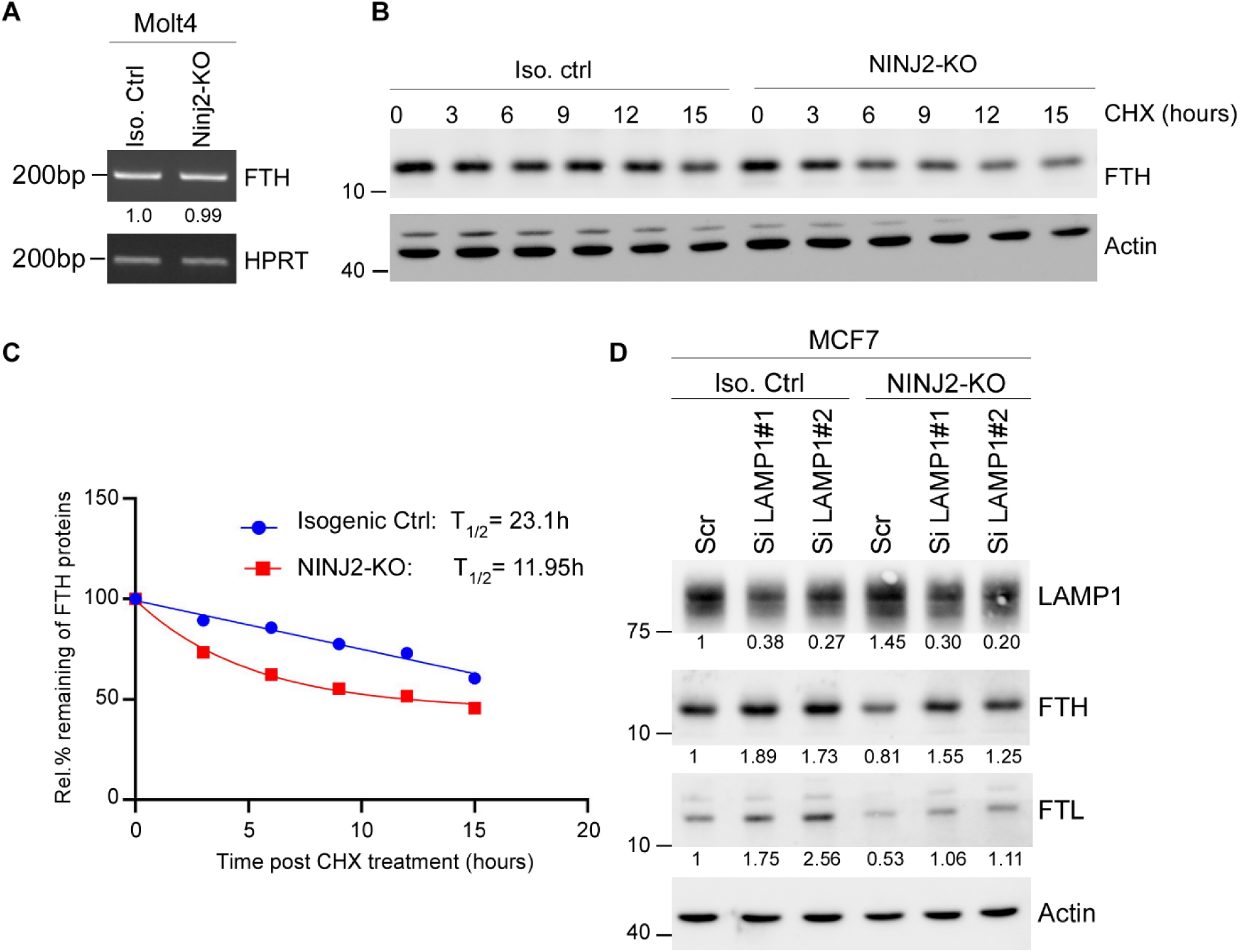
Loss of NINJ2 promotes ferritin degradation. (**A**) The level of FTH and HPRT transcripts was measured in isogenic control and NINJ2-KO Molt4 cells. The relative level of FTH transcripts in control cells was arbitrarily set as 1.0 and the relative fold change of transcripts was shown below each lane. (**B**) Isogenic control and NINJ2-KO MCF7 cells were treated with cycloheximide (50 μg/mL from 3 to 15 hours. The cell lysates were collected and subjected to western blot analysis using FTH and actin antibodies. (**C**) The level of FTH and actin protein in (B) were quantitated and the relative protein half-life of FTH was calculated by GraphPad Prism software. (**D**) Isogenic control and NINJ2-KO MCF7 cells were transiently transfected with scrambled siRNA or siRNAs against LAMP1 for 3 days, followed by western blot analysis with antibodies against LAMP1, FTH, FTL, and actin. The relative level of LAMP1, FTH, and FTL protein in control cells was arbitrarily set as 1.0 and the relative fold change of transcripts was shown below each lane.

### Loss of NINJ2 promotes ferroptosis

Recent studies have shown that LMP releases reactive iron into the cytosol and promotes lipid peroxidation, thereby serving as a critical initiating event in ferroptosis, an iron-dependent form of programmed cell death [33, 38]. Since loss of NINJ2 increases the level of labile iron (Fig. 3A-B), we thus examined whether NINJ2 modulates ferroptosis. As FTH serves as a protector against ferroptosis, its expression was first examined in isogenic control and NINJ2-KO cells treated with or without RSL3 and Erastin, both of which are ferroptosis inducers [39–41]. Indeed, we found that the level of FTH was decreased by NINJ2-KO regardless of RSL3 or Erastin treatment in both Molt4 and MCF7 cells (Fig. 5A-B), suggesting that NINJ2-deficiency promotes ferroptosis. To further test this, isogenic control and NINJ2-KO Molt4 cells were treated with various doses of RSL3 or Erastin, and cell viability was assessed. We found that loss of NINJ2 significantly sensitized Molt4 cells to RSL3 treatment, reducing the IC_50_ from 0.31 μM in isogenic control cells to 0.16 μM in NINJ2-KO cells (Fig. 5C). Similarly, NINJ2 deficiency enhanced the sensitivity of Molt4 cells to Erastin, as evidenced by decreased IC₅₀ in NINJ2-KO cells (Fig. 5D). To validate these findings, isogenic control and NINJ2-KO MCF7 cells were treated with RSL3 or Erastin. Consistently, we found that loss of NINJ2 promoted ferroptosis in MCF7 cells, as indicated by a marked reduction in their IC₅₀ values (Fig. 5E-F). To further verify this, colony formation assay was performed. We found that under mock treatment conditions, NINJ2-KO suppressed colony formation, consistent with previous reports [13]. Importantly, RSL3 and Erastin markedly reduced colony formation in MCF7 cells, which was further inhibited by NINJ2-KO (Fig. 5G-H).

**Figure 5.**
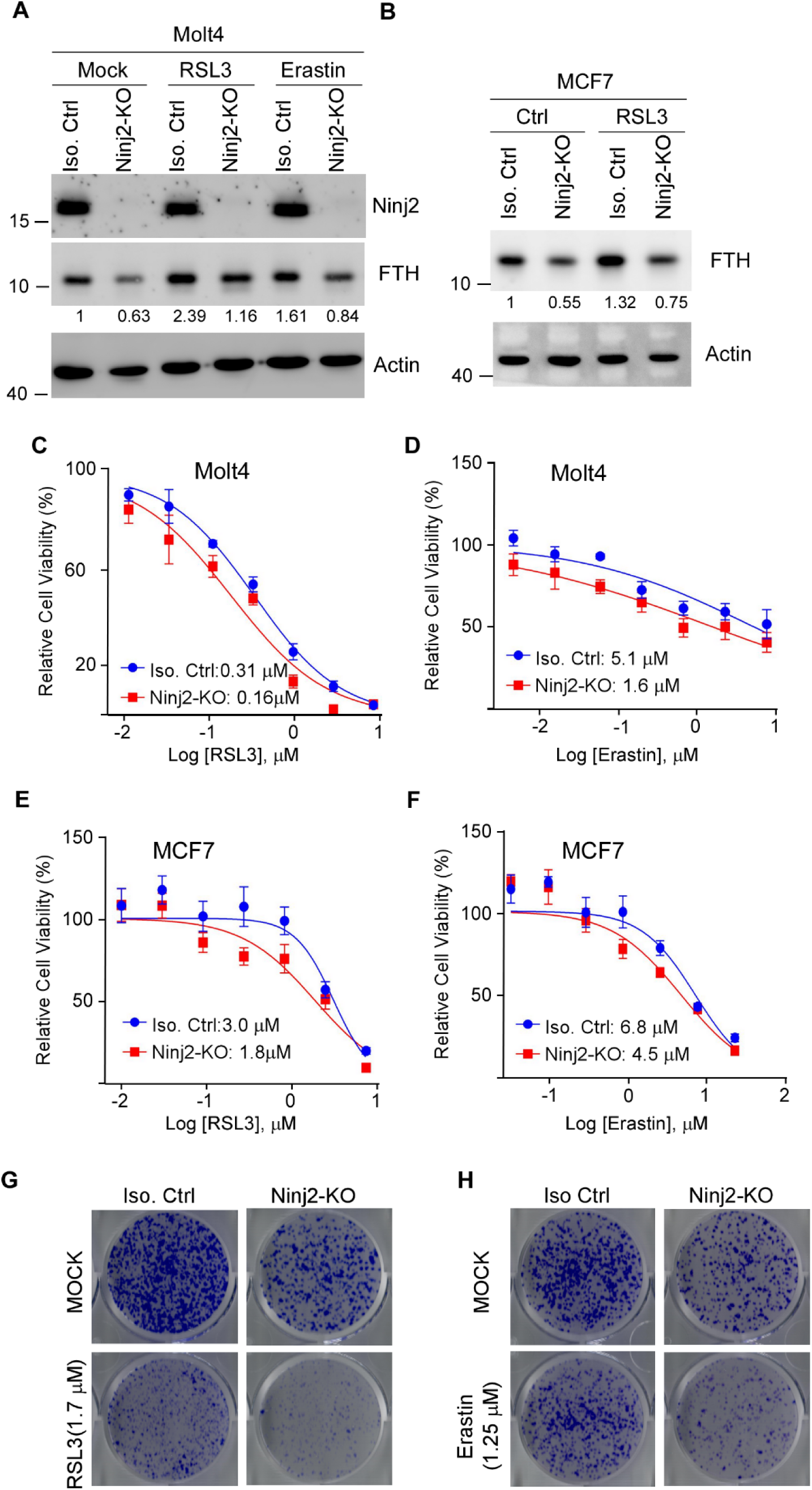
Loss of NINJ2 promotes ferroptosis. **(A**) Isogenic control and NINJ2-KO Molt4 cells were treated with RSL3 (1.25 μM) or Erastin (2.5 μM) for 8 hours, and the level of Ninj2, FTH, and actin were measured by western blot analysis. The relative protein level of FTH in control cells were arbitrarily set as 1.0 and the relative fold change was shown in below each lane. (**B**) Isogenic control and NINJ2-KO MCF7 cells were treated with RSL3 (1.25 μM) for 8 hours, followed by western blot analysis to detect FTH, and actin. The relative protein level of FTH in control cells were arbitrarily set as 1.0 and the relative fold change was shown in below each lane. (**C-D**) Isogenic control and NINJ2-KO Molt4 cells were treated with RSL3 (C) or Erastin (D) from 0-7.29 μM for 48 hours. The relative cell viability was measured using CellTiter-Glo Viability Assay kit. The relative Cell viability is calculated as a percentage of untreated (control) cells. Data points represent the mean ± SD from four representative experiments. Curves were fitted using nonlinear regression using GraphPad Prism Software. IC50 values represent the drug concentration required to achieve 50% inhibition of maximal proliferation capacity. (**E-F**) Isogenic control and NINJ2-KO MCF7 cells were treated with RSL3 (E) or Erastin (F) from 0-7.29 μM for 48 hours, followed by cell viability assay. The IC50 was calculated with nonlinear regression analysis using GraphPad Prism Software. (**G-H**) Colony formation was performed with isogenic control and NINJ2-KO MCF7 cells were treated with or without RSL3 or Erastin for 24 hours. The drugs were then withdrawn to allow colonies to grow for 3 weeks.

### NINJ2, FTH1 and FTL are overexpressed in hepatocellular and breast carcinomas and are positively associated among one another

Previous reports have shown that Ferritins promotes tumorigenesis through regulating iron homeostasis and oxidative stress as well as suppressing ferroptosis. To determine whether the NINJ2-Ferrrtins axis contributes to cancer development, we searched the TCGA database for the association between NINJ2 and ferritin in several types of human cancers. Interestingly, we found that NINJ2, FTH, and FTL were all up-regulated in hepatocellular carcinoma (Fig. 6A-C) and breast cancer (Fig. 6D-F), both of which are iron-addicted malignancies. Further, we found that NINJ2 expression was positively associated with FTH and FTL in both hepatocellular carcinoma (Fig. 6G-H) and breast carcinoma (Fig. 6I-J). Together, these findings suggest that NINJ2 may function in concert with ferritin to promote tumor cell growth especially in iron-addicted tumors, which may represent a therapeutic vulnerability for these types of cancers.

**Figure 6.**
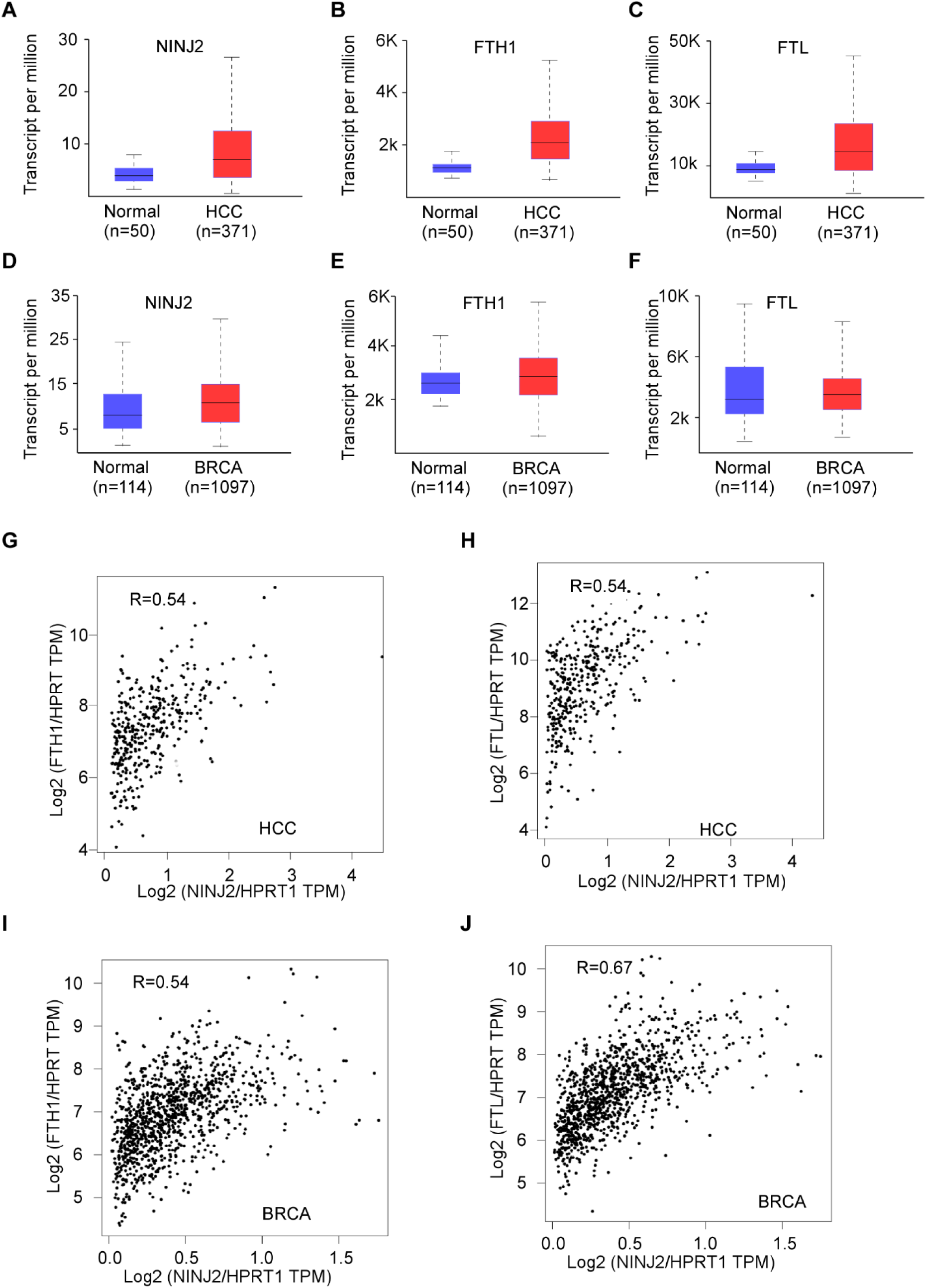
NINJ2 and Ferritins are overexpressed in hepatocellular and breast carcinomas and are positively associated with one another. **(A-C)** Boxplot shows the relative expression of NINJ2 (A), FTH (B), and FTL (C) in normal and hepatocellular carcinomas. The analysis was performed using UACLAN database. (**D-F**) Boxplot shows the relative expression of NINJ2 (A), FTH (B), and FTL (C) in normal and breast carcinomas. The analysis was performed using UACLAN database. (**G**) NINJ2 expression is positively associated with FTH (G) in hepatocellular carcinomas. The analysis was performed using the GEPIA2 database (http://gepia2.cancer-pku.cn/#correlation). Statistical analysis suggests a strong correlation between NINJ2 and FTH expression in hepatocellular carcinomas (Pearson’s *r* =0.54). (**H**) NINJ2 expression is positively associated with FTL (H) in hepatocellular carcinomas (Pearson’s *r* =0.54). (**I-J**) NINJ2 expression is positively associated with FTH (I, Pearson’s *r* =0.54) and FTL (J, Pearson’s *r* =0.67) in breast carcinomas.

## Discussion

NINJ2 is a multifaceted protein and participates in various biological and pathological processes. However, the underlying mechanisms underlying these processes are not fully understood. Here, we identify NINJ2 as a critical regulator of lysosomal integrity and ferroptosis. Specifically, we demonstrate that NINJ2 localizes to lysosomes and interacts with LAMP1, and that loss of NINJ2 enhances LMP. We also found that loss of NINJ2 increases the level of cytosolic labile iron by promoting lysosomal ferritin degradation. Consequently, NINJ2 deficiency sensitizes cells to ferroptosis. Finally, we showed that NINJ2 and ferritin are co-overexpressed and positively correlated in several iron-addicted cancers, including hepatocellular carcinoma and breast cancer, suggesting that the NINJ2-FTH axis may be targeted for these types of cancer. These data let us speculate that loss of NINJ2 leads to low-grade lysosomal membrane damage, leading to leakage of labile iron into the cytosol. The released iron increases oxidative stress and promotes ferritin degradation in lysosomes, which further elevates cytosolic iron levels, creating a feed-forward loop that sensitizes cells to ferroptosis. A model to elucidate the role of NINJ2 in maintaining lysosomal membrane integrity and iron homeostasis is proposed in Figure 7.

**Figure 7.**
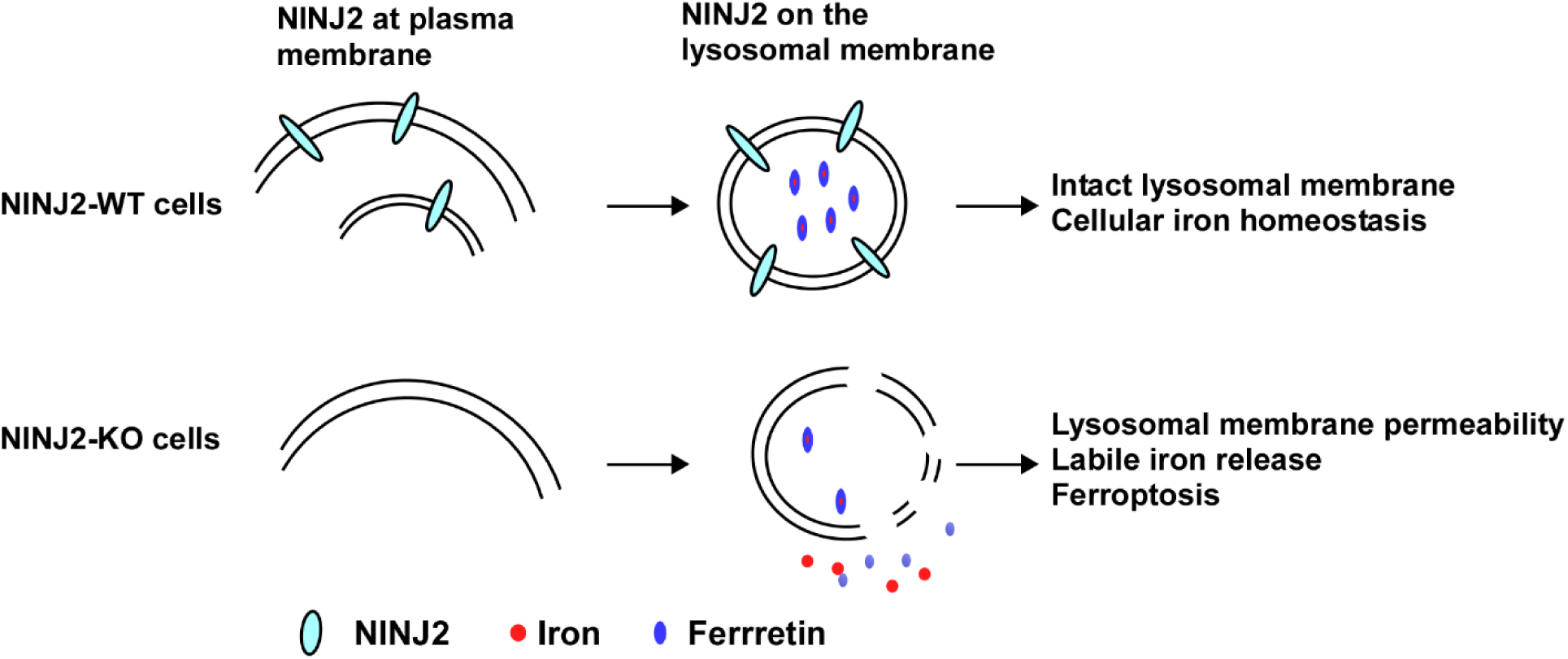
A model to elucidate the role of NINJ2 in maintaining lysosomal membrane integrity and iron homeostasis.

Another interesting observation is that NINJ2 is required for Ferritin expression. Our data indicated the NINJ2-deficiency leads to reduced expression of Ferritin (Fig. 3C-F). Moreover, we found that reduced Ferritin expression by loss of NINJ2 is due to enhanced lysosomal degradation (Fig. 4B-C). In support of this, we showed that knockdown of LAMP1 reverses the inhibition of ferritin expression observed in NINJ2-knockout cells (Fig. 4D). Although not yet experimentally confirmed, we tentatively propose that the enhanced degradation of ferritin in NINJ2-KO cells results from increased labile iron in these cells owing to the low-grade LMP. On the other hand, the NINJ2-Ferritin axis may be explored as viable strategies for cancer management. In support of this notion, we found that NINJ2 is positively associated with Ferritins in iron-addicted cancers, such as HCC and Breast cancer (Fig. 6). Notably, most iron-addicted cancers rely on intact lysosomal function to shield excess iron and evade oxidative stress. Thus, targeting NINJ2 may unmask a convergent vulnerability, rendering tumor cells more susceptible to ferroptosis. Notably, we have developed a small peptide derived from N-terminal extracellular domain that can enhance pyroptosis [42]. It would be interesting to determine whether this peptide can enhance ferroptosis.

Recent studies have shown that lysosomal leakage does not always lead to cell death since minor lysosomal membrane damage can be repaired via the ESCRT complexes [21, 43]. Interestingly, low-grade lysosomal leakage can lead to limited release of lysosomal contents, such as iron and acid hydrolases, thereby influencing diverse cellular processes such as inflammatory responses [44]. The finding that loss of NINJ2 leads to low-grade LMP (Fig. 2), is consistent with previous observation that loss of NINJ2 promotes pyroptosis and inflammation [9]. How does loss of NINJ2 lead to LMP? One possibility is that loss of NINJ2 may alter membrane fluidity through lipid remodeling. Indeed, we showed previously that loss of NINJ2 alters lipid metabolism and NINJ2-deficiency leads to marked increase in ceramide level [9]. Notably, ceramide is a critical lipid mediator that destabilizes lysosomal membranes, acting as a potent trigger for LMP and subsequent cell death. Another possibility is through forming a complex with LAMP1 (Fig. 1B-E). We postulate that the NINJ2-LAMP1 complex acts as a protective scaffold on the lysosome surface and thereby prevents the leakage of lysosomal content. Interestingly, we also observed that NINJ2-KO leads to increased LAMP1 expression at both transcript and protein levels (Fig. 2 B-E and Supplemental Figure 1), which may be a compensatory protective response. Indeed, elevated LAMP1 levels have been found to stabilize damaged lysosomal membranes and facilitate lysosomal repair, thereby preventing cells from death induced by membrane rupture [45]. LAMP1 transcript is known to be updated by TFEB (Transcription Factor EB), a master regulator of the lysosomal-autophagy system [46]. When LMP occurs, TFEB can translocate to nuclear and activate several lysosome-related genes, including LAMP1. Notably, phosphorylation of TFEB by mTORC1 at Serine 211 will inactivate TFEB by preventing its translocation to nuclear [47]. Thus, it would be interesting to determine whether NINJ2 plays a role in modulating TFEB phosphorylation. In addition, the tumor suppressor p53 has been reported to engage in intricate crosstalk with TFEB to regulate basal autophagy [48]. Interestingly, p53 expression is found to be increased by NINJ2-KO [13]. It is therefore of interest to determine whether NINJ2 regulates TFEB activity through p53 or modulates the p53-TFEB signaling axis to maintain lysosomal integrity. Finally, NINJ1, the other member of the Ninjurin family, has been shown to promote pyroptosis. Our previous studies demonstrated that NINJ2 may antagonize NINJ1 by forming a complex with NINJ1. It will therefore be of considerable interest to determine whether NINJ1 also regulates LMP and whether it exhibits a role opposite to that of NINJ2.

Overall, our data expand the functional landscape of NINJ2 and establish it as a potential therapeutic node at the intersection of lysosomal integrity, iron metabolism, and ferroptosis.

## Materials and Methods

### Reagents

Anti-FTH (Cat #4393S), anti-LAMP1 (Cat #3243S and Cat #9091S), and anti-GAPDH (Cat #2118L) were purchased from Cell Signaling Technology. Anti-actin (Cat #sc-8432) and anti-Galectin 3(Cat #sc-32790) were purchased from Santa Cruz Technology. Anti-Flag (Cat# 80801-2-RR) and anti-FTL (Cat #10727-1-AP) were purchased from Proteintech. Anti-NINJ2 was custom-made as previously described [49] and affinity purified. Goat anti-Rabbit IgG (FITC) (Cat #ab6717) and Goat Anti-Mouse IgG (FITC) were purchased from abcam. Goat anti-Mouse IgG (Alexa 555) and Goat anti-Rabbit IgG (Alexa 555) were purchased from Life technology. Proteinase inhibitor cocktail (Cat #78429), Trizol Reagent (Cat #15596026), RNAiMAX (Cat #13778), and RevertAid First Strand cDNA Synthesis Kit (Cat #K1621) were purchased from Life Technologies. The WesternBright Sirius HRP substrate (Cat #K12043-D20) was purchased from Advansta. JetPRIME transfection reagent (Cat #101000046) was purchased from Polyplus. Protein A/G magnetic beads (Cat # HY-K0202), RSL3 (Cat #HY-100218A), and Erastin (Cat # HY-15763) were purchased from MedChemExpress. QuantiChrom™ Iron Assay Kit (Cat # DIFE-250) was purchased from BioAssay Systems

### Cell culture

MCF7, Molt 4 and 293T cells were purchased from the American Type Culture Collection (ATCC). Since all cell lines from ATCC have been thoroughly tested and authenticated, we did not authenticate the cell lines used in this study. MCF7 and 293T were cultured in Dulbecco’s modified Eagle’s medium (Gibco, Cat #12100061) supplemented with 10% fetal bovine serum (Gibco, Cat # 10437-028) and Penicillin-Streptomycin (Gibco, Cat #15140122). Molt4 cells were cultured in RPMI 1640 medium (Cat# 11875093) supplemented with 10% fetal bovine serum and Penicillin-Streptomycin. Isogenic control and NINJ2-KO MCF7 and Molt4 cells were previously generated [9, 13]. Briefly, MCF7 and MOLT-4 cells were transfected with two gRNAs targeting NINJ2 and selected with puromycin for 3 weeks. Individual clones were then isolated, and NINJ2 knockout was confirmed by Western blot analysis. All the cells were used below passage 25 or within 2 months after thawing.

### Immunofluorescence mirocoscopy

MCF7 cells expressing 3 x Flag-tagged NINJ2 were fixed with 3.7% formaldehyde in phosphate buffered saline (PBS), permeabilized with 0.2% Triton X-100 in PBS, and blocked with 2% bovine serum albumin in PBS. The cells were then stained with primary antibodies, followed by fluorophore-conjugated secondary antibodies. The cells were mounted with ProLong Gold with DAPI and observed with Leica SP8 confocal microscope with a 40x oil immersion objective or x63 oil immersion objective. The dilutions of the primary antibodies were 1:100 for anti-LAMP1, 1:100 for anti-Ninj2, and 1:100 for anti-Gal3. The dilutions for secondary antibody were 1:500 for Alexa 555-conjugated and 1:1000 for FITC-conjugated.

### Lysosomal staining

Cells were incubated with CytoFix Red Lysosomal stain (AAT, Cat# 23210) for 30 minutes at 37℃. The cells were fixed, permeabilized, and blocked, and then stained with primary antibodies followed by fluorophore-conjugated secondary antibodies as described above. The dilution for CytoFix Red was 1:500.

### Proximity ligation assay (PLA assay)

PLA assay was carried out by using DuoLink PLA assay kit (Millipore Sigma, Cat# DUO92008, DUO92001, and DUO92005) according to the manufacturer’s protocol. Briefly, cells were fixed with 3.7% formaldehyde in PBS, permeabilized with 0.2% Triton X-100 in PBS, and blocked using the blocking reagent provided in the kit. The cells were then incubated with primary antibodies at 4 °C overnight (∼20 h). On the following day, cells were incubated with the PLUS and MINUS probes at 37 °C for 1 h, followed by incubation with DNA ligase at 37 °C for 30 min and subsequent incubation with DNA polymerase together with the fluorescent probe at 37 °C for 100 min. The cells were mounted with ProLong Gold with DAPI and observed with Leica SP8 confocal microscope as described above.

### Transient transfection

SiRNA transfection was performed with RNAiMAX according to user’s manual. The sequence for scrambled siRNA was 5’-GCA GUG UCU CCA CGU ACU A-3’. The sequences for LAMP1 siRNA were 5’-CAG CAA UGU UUA UGG UGA AUU-3’, and 5’-CCA AAG AAA UCA AGA CUG UUU-3’.

### Western Blot and Immunoprecipitation (IP) analyses

Western blot analysis was performed as previously described [50]. Briefly, whole-cell lysates were resolved on 8-13% SDS–polyacrylamide gels and transferred to nitrocellulose membranes. Membranes were incubated with primary and secondary antibodies, followed by detection using enhanced chemiluminescence and visualization with VisionWorks LS software (version 8.0; Analytik Jena, Jena, Germany). For IP analysis, Cells were lysed in an IP lysis buffer (1% NP-40, 50 mM Tris-HCl (pH 8.0), 150 mM NaCl, and 1 mM EDTA) supplemented with proteinase inhibitor cocktail. The cell lysates with incubated with 1 μg of primary antibody and protein A/G magnetic beads at 4 ℃ overnight and the immunocomplex were subjected to western blot analyses to detect protein-protein association.

### Cell viability

4×10^3 Cells were seeded per well in 96-well plates (100 μl per well) in quadruplicate and allowed to adhere overnight. Next day, cells were treated with varying concentrations of RSL3 or Erastin (0-7.29 μM) for 48 hours. Cell viability was assessed by adding 100 μl of CellTiter-Glo reagent (Promega) to each well and incubated for 10-minute at room temperature. Following incubation, luminescence was measured using a SpectraMAX Gemini Microplate Reader (Molecular Devices, Silicon Valley, CA, USA). The viability in the control group was set as 100% and the relative cell viability was calculated as a percentage of treatment group vs control group. The IC50 was calculated using GraphPad Prism 10.

### Colony formation assay

2×10^3 Cells were seeded per well in a 6-well plate in triplicate. At 48 hours, cells were treated with various amounts of RSL3 or Erastin for 24 hours and the drug was withdrawn. Cells were then cultured in regular medium for 2 weeks to allow colonies to form. The colonies were then fixed with methanol/glacial acetic acid (7:1) and stained with 0.1% of crystal violet at room temperature.

### Total RNA isolation and RT-PCR analysis

Total RNA was isolated with Trizol reagent as described according to the user’s manual (Life Technologies). 3 μg of total RNA was used to synthesize cDNA by using RevertAid First Strand cDNA Synthesis Kit, followed by PCR analysis. The program used for amplification was (i) 94 °C for 5 min, (ii) 94 °C for 45 s, (iii) 58 °C for 45 s, (iv) 72 °C for 30 s, and (v) 72 °C for 10 min. From steps 2 to 4, the cycle was repeated 28–35 times depending on the targets or 22 times for actin and GAPDH. The primers for LAMP1 were a forward primer, 5′-AGG ACA TAC ACT CAC TCT C-3′, and a reverse primer, 5′-GTG CCA CTA ACA CAT CTG-3′. The primers for FTH were a forward primer, 5′-CGA TGA TGT GGC TTT GAA GA-3’, and a reverse primer, 5′-AAT GGG GGT CAT TTT TGT CA-3′. The primers for HPRT1 were a forward primer, 5′-TAT GGC GAC CCG CAG CCC T-3′, and a reverse primer, 5′-CAT CTC GAG CAA GAC GTT CAG-3′. The primers for Actin were a forward primer, 5′-CTG AAG TAC CCC ATC GAG CAC GGC A-3′, and reverse primer, 5′-GGA TAG CAC AGC CTG GAT AGC AAC G-3′.

### Labile iron measurements

Cells were treated with ferric ammonium citrate (300 μg/ml) for 16 hours. After treatment, 1×10^5^ cells were collected and lysed in 200 μl of RIPA buffer. Cell lysates were then subjected to labile iron assay using the QuantiChrom Iron Assay Kit according to manufacturer’s instructions.

### Protein half-life measurement

Cells were mock-treated or treated with cycloheximide (50 μg/ml) from 0-15 hours. Cell lysates collected at each timepoint were subjected to western blot analysis to detect FTH and actin. Band intensities at the different time points were quantified using the VisionWorks LS software, normalized to actin and plotted in graph as relative percentage of remaining protein.

### Statistical analysis

Student’s t test was used for statistical analysis. p<0.05 is considered as significant.

## Acknowledgement

This work was supported in part by National Institutes of Health R01 grants (CA272753), a UC Davis Cancer Center Core Support Grant (CA093373).

## Data Availability

All study data are included within the article.

## Author Contributions

J.Z. and X.C. designed the research. J.Z., M.B., K.N., and Y.S. performed the research. J.Z., M.B., K.N., and Y.S.and X.C. analyzed the data. J.Z. and X.C. wrote the paper. All authors have read and agreed to the published version of the manuscript.

---

**Supplemental Figure 1.**
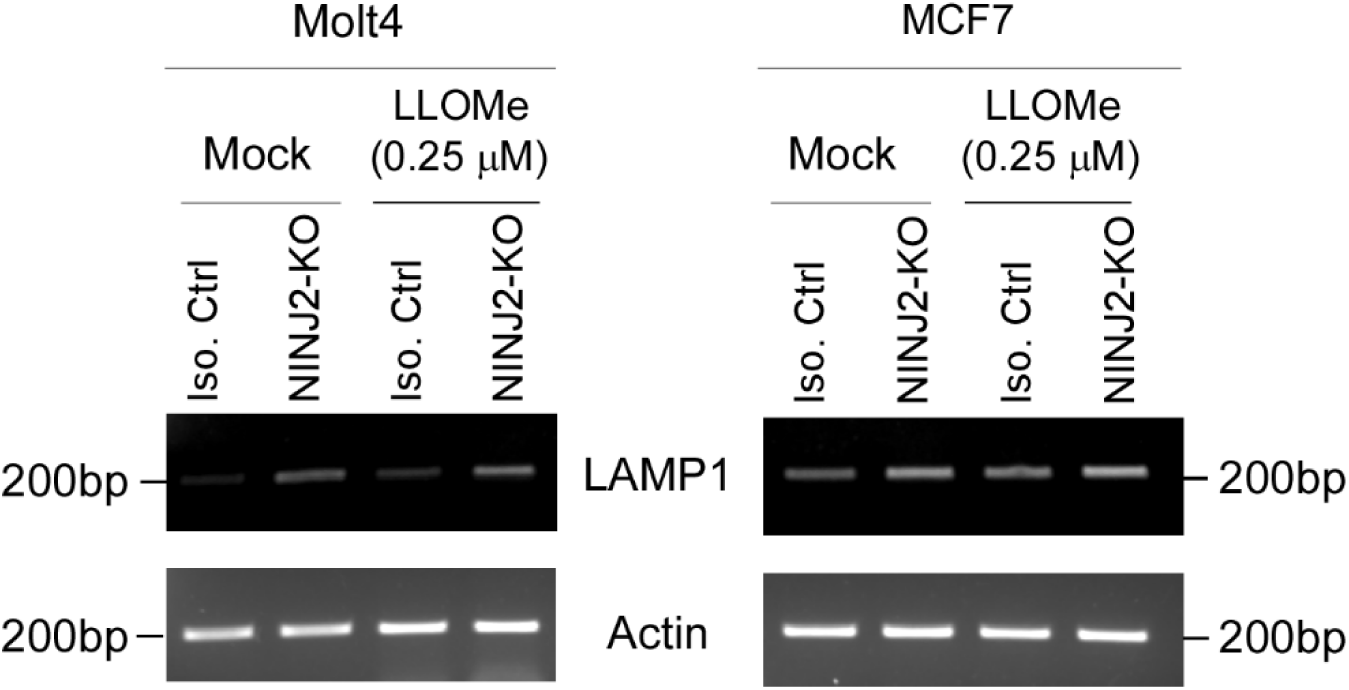

## Notes

### Competing Interest Statement

The authors have declared no competing interest.

### Summary of Updates

The manuscript has been peer-reviewed and a revision is made to address reviewer's conerns.

