## Supplementary figures and images for "Nerve Injury-Induced Protein 2 preserves lysosomal membrane integrity to suppress ferroptosis"

### Supplemental Figure 1

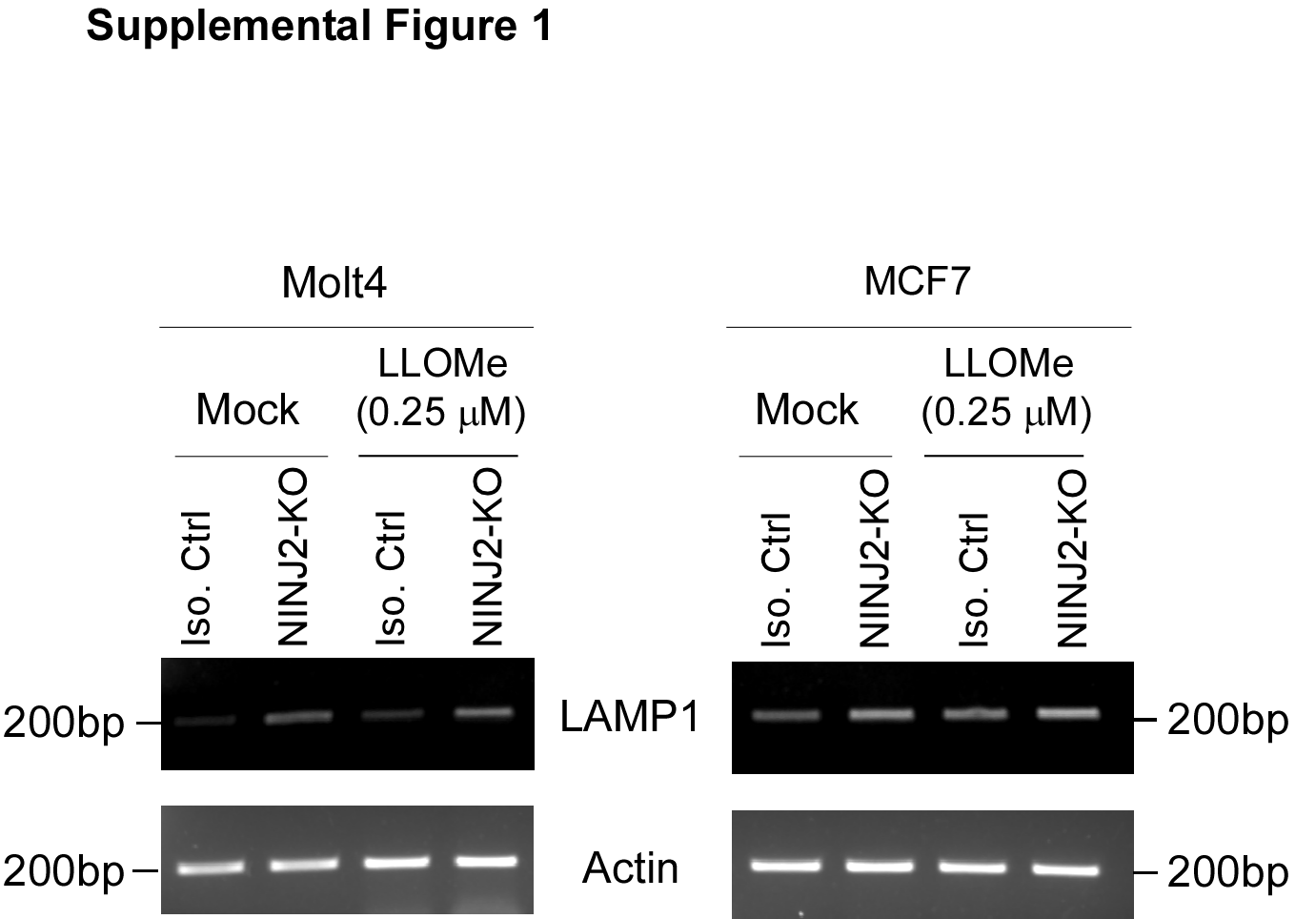
